# Microbially assisted rescue of host populations

**DOI:** 10.64898/2026.10.15.759220

**Authors:** Bob Week, Hildegard Uecker, Hinrich Schulenburg

## Abstract

Rapid environmental change can drive populations into decline but host genomes may adapt in time to rescue the population from extinction. Host-associated microbes may alter this race. We here develop a population-genetics model to characterize the dynamics and consequences of microbe-assisted host population rescue. In our model, we replace one locus of a classical two-locus system with microbial carriage. Under weak selection the microbial variables settle to a quasi-microbial equilibrium (QME) conditional on the host allele, reducing the coupled dynamics to host allele frequency and abundance. The reduction identifies two microbial contributions that shape rescue. Additive benefits of microbial carriage on host fitness raise host growth immediately without affecting evolution and lift the floor of the demographic rescue-U. By contrast, positive interactions between the rescue allele and the microbe accelerate evolution and narrow the rescue-U instead of buffering initial decline. A microbe can therefore buy time, speed adaptation, or be required for recovery when neither contribution suffices alone. Imperfect microbial inheritance, however, can impede microbe-assisted rescue. When selection acts through fecundity and carriers transmit the microbe to only some offspring, adaptation generates non-carrying descendants faster than transmission replaces them. When that loss is sufficiently strong, recovery can fail even though the allele, the microbe, and their interaction are each individually beneficial. We call this selection-amplified dilution. Together, our model integrates evolutionary rescue theory with multilocus population genetics and evolutionary epidemiology to define a framework of microbe-assisted host rescue, thereby identifying key factors driving or compromising the rescue process.

**Significance:** Microbes living in and on animals and plants influence key components of host fitness, raising the question of whether they can determine if a population survives severe environmental change. We here develop a mathematical model that separates two ways they do it. A microbe can raise host growth directly, cushioning the population while its genes catch up, or it can influence how strongly a beneficial host gene is favored, speeding up evolutionary adaptation. The model additionally reveals an unexpected failure mode: when successful hosts pass the microbe to only some of their offspring, adaptation itself can erode a beneficial microbe enough to prevent recovery. Overall, our model identifies distinct mechanisms and their signatures of how associated microbes can enhance evolutionary rescue of their hosts.

---

Rapid environmental change can push populations to the brink of extinction, but genetic adaptation may restore growth, a phenomenon termed evolutionary rescue. Models of evolutionary rescue therefore ask not only whether beneficial alleles spread, but whether they spread fast enough to prevent extinction [1, 2]. This problem is usually framed as adaptation by the host genome. Yet, hosts also acquire fitness-affecting microbes through vertical inheritance, social transmission, and environmental acquisition [3–8]. These microbes can alter host performance immediately and persist across host generations [9]. They therefore create an additional route through which a declining population may delay extinction or recover entirely. Existing theory has developed frameworks for quantifying microbiome contributions to heritable host variation and predicting microbial responses to host-level selection [10–17]. What remains unclear is when and how these microbial effects on host traits translate into population recovery. In the setting we consider, where genetic adaptation proceeds through a beneficial host allele, microbes may alter rescue through two distinct routes: by slowing demographic decline or by accelerating the spread of the beneficial allele. The central question is therefore not simply whether microbes are beneficial, but how microbial transmission and host–microbe interaction partition their effect between demographic buffering and genetic adaptation. To date, the exact dynamics and the resulting consequences of such microbe-mediated host rescue are largely unexplored, even though they are likely of key importance for conservation biology in light of the ubiquity of host-associated beneficial microbes.

To address this current knowledge gap, we here develop a mathematical model that simplifies the complexity of host– microbiome assemblages and incorporates microbe presence as a binary host trait responding to horizontal acquisition, clearance, and imperfect inheritance. We define a four-state host–microbe model that preserves the logic of a classical two-locus system while replacing one locus with microbial carriage. The same architecture also describes conjugative plasmids, in which accessory genes spread horizontally and are lost through imperfect segregation. We introduce a weak-selection quasi-microbial equilibrium (QME), analogous to quasi-linkage equilibrium [18–20], and use it to reduce the dynamical system to two equations – for the the host allele and population abundance. The reduction yields two first-order results. The microbial selection coefficient, *s*_*m*_, buffers demographic decline in proportion to baseline microbial prevalence. The non-additive gene–microbe interaction coefficient, *e*, changes the rate at which the host rescue allele spreads. The same reduction exposes a countervailing mechanism: when selection acts through fecundity and microbial inheritance is incomplete, adaptation can dilute a beneficial microbe and reverse host–microbe associations. We call this selection-amplified dilution. It can deepen demographic decline or prevent rescue even when the host allele, the microbe, and their interaction are all individually beneficial.

Below, we first formulate the host–microbe model and derive the QME reduction, then use it to separate demographic buffering from evolutionary acceleration and to identify the conditions for selection-amplified dilution. We then discuss connections to existing theory, empirical tests, and biological extensions, with derivations and numerical validation provided in the Materials and Methods and the SI Appendix.

## Results

### A model of microbially assisted host rescue

Let *p*_*g*_ be the marginal frequency of hosts carrying the rescue allele and *p*_*m*_ the marginal frequency carrying a focal microbe. For simplicity, hosts are haploid, and the rescue allele is transmitted faithfully from parent to offspring. If *p*_*gm*_ is the frequency carrying both, their host–microbe association is (Fig. 1A)

**Figure 1:**
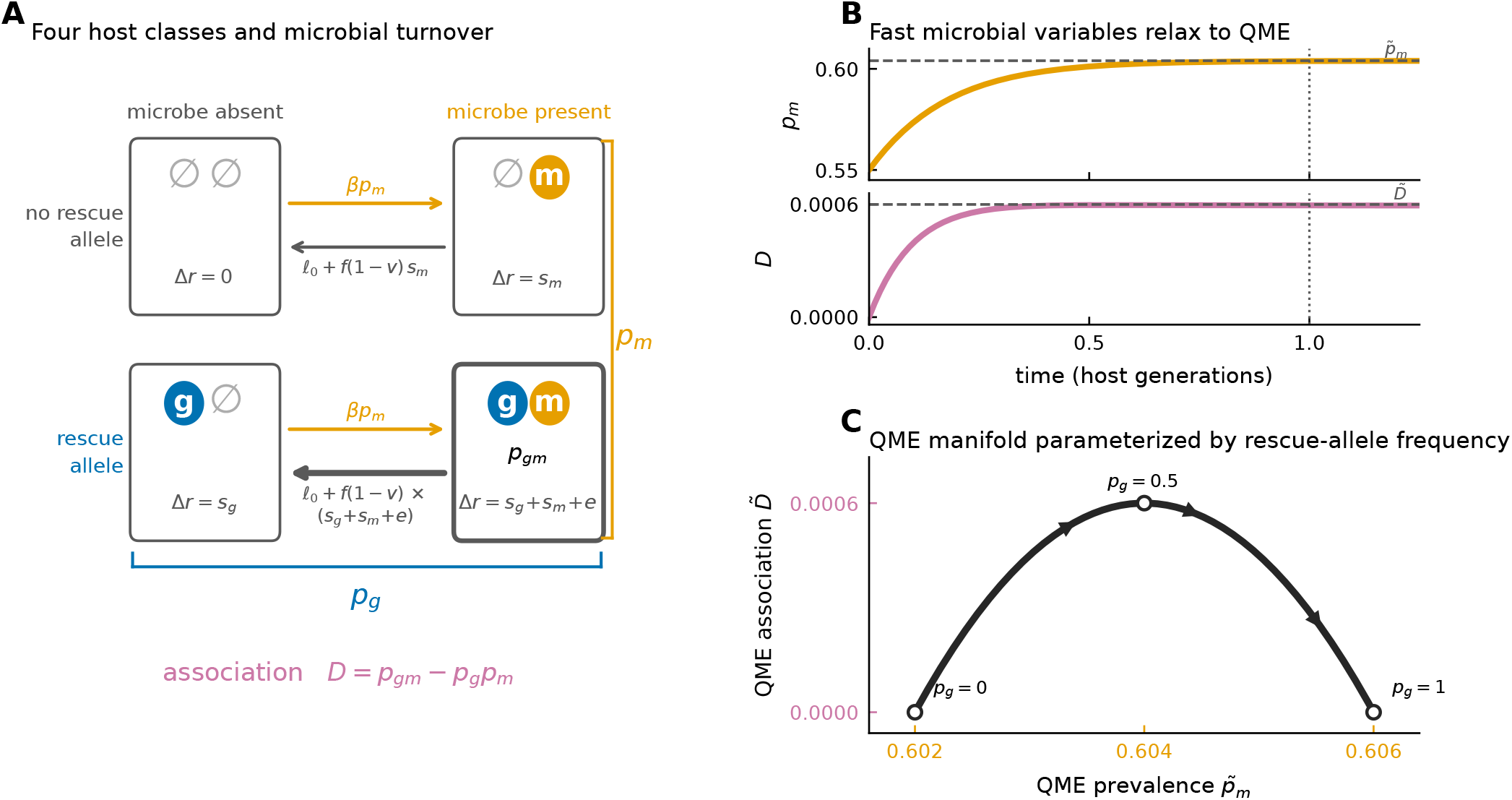
Host–microbe model and quasi-microbial equilibrium (QME) reduction. (*A*) The four host classes are determined by presence or absence of the rescue allele and microbe. Here Δ*r* denotes the class-specific change in Malthusian fitness relative to hosts carrying neither element; the double-carrier effect *s*_*g*_ + *s*_*m*_ + *e* therefore includes the non-additive excess *e*. Non-carriers acquire the microbe at rate *βp*_*m*_, whereas microbial loss combines baseline loss *ℓ*_0_ with the fecundity-mediated failed-inheritance term *f* (1 *− v*)Δ*r*. Marginal rescue-allele frequency *p*_*g*_, microbial prevalence *p*_*m*_, and their association *D* = *p*_*gm*_ *− p*_*g*_*p*_*m*_ follow from these four classes. (*B*) At fixed *p*_*g*_ = 0.5, microbial prevalence *p*_*m*_ and host–microbe association *D* relax toward their QME values 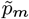 and 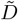 (dashed lines); the vertical dotted line marks one host generation. (*C*) The QME values 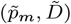 form a manifold parameterized by *p*_*g*_, with marked positions at *p*_*g*_ = 0, 0.5, and 1. Panels B–C use *β* = 10, *ℓ*_0_ = 4, *ϕ* = 0, *s*_*g*_ = 0.03, *s*_*m*_ = 0.05, and *e* = 0.10, giving 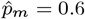; panel B starts from *p*_*m*_(0) = 0.55 and *D*(0) = 0.

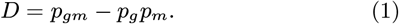

Thus *D* = 0 corresponds to random association, whereas *D >* 0 means that the allele and microbe co-occur more often than expected from their marginal frequencies. Let *s*_*g*_ and *s*_*m*_ denote the selection coefficients for the rescue allele and microbial carriage. Their joint Malthusian fitness effect is *s*_*g*_ + *s*_*m*_ + *e*, where *e* is the non-additive gene–microbe interaction coefficient. We focus on *s*_*m*_ ≥ 0 and *s*_*m*_ + *e >* 0, so microbial carriage is non-deleterious without the rescue allele and beneficial in its presence. The full model tracks the four combinations of rescue-allele and microbial carriage illustrated in Fig. 1A; their class-specific dynamics and reduction to (*p*_*m*_, *p*_*g*_, *D*) are given in SI Appendix, Section S1.

Microbial acquisition is modeled as frequency dependent: each non-carrier acquires the microbe at rate *βp*_*m*_, where *β* is the horizontal transmission coefficient. A fraction 1 − *v* of a carrier’s offspring lack the microbe, and in addition, carriers are cleared at rate *γ*. We write *ℓ*_0_ = *b*(1 − *v*) + *γ* for baseline loss through failed vertical inheritance and clearance, where *b* is the fecundity of individuals that carry neither the rescue allele nor the microbe. A fraction *f* of selection acts through fecundity and 1 − *f* through survival, so selection-dependent failed inheritance scales with *f* (1 − *v*). Common density-dependent mortality *cN* only bounds host abundance and cancels from the frequency equations. After environmental change,

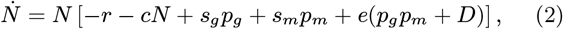

with *r >* 0, so hosts carrying neither element have baseline decline rate *r* at low density; at finite abundance, competition contributes the additional term −*cN* to per-capita growth. The full dynamics and their continuous-time Markov-chain foundation are given in SI Appendix, Sections S1–S2.

### The quasi-microbial equilibrium

Without selection at the host level (i.e., *s*_*m*_ = *s*_*g*_ = *e* = 0), microbial carriage approaches

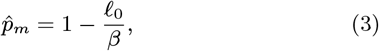

when *β > ℓ*_0_. A ‘hat’ denotes this selection-free endemic equilibrium. We next turn to the quasi-microbial equilibrium, denoted by ‘tildes’ (see SI Appendix, Section S3 for a full derivation). Intuitively, at fixed rescue-allele frequencies *p*_*g*_, the microbial frequency *p*_*m*_ and the allele-microbe association *D* rapidly relax toward limiting values (their quasi-microbial equilibrium, Fig. 1B); under weak selection (*s*_*g*_, *s*_*m*_, *e* ≪ 1), they therefore track the slower dynamics of *p*_*g*_. To first order,

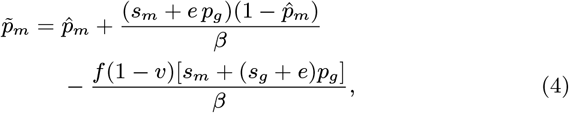

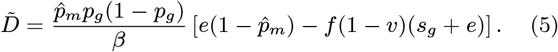

As *p*_*g*_ changes, these QME values define the manifold shown in Fig. 1C. The first correction to 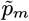 with regard to 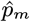 is selection for microbial carriers; the second is due to selection-amplified dilution through births that fail to inherit the microbe. Substituting the QME (Eqs. 4–5) into the exact host allele equation (S1b) gives

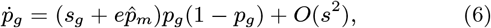

where *s* denotes the scale of *s*_*g*_, *s*_*m*_, and *e*. Host abundance changes according to

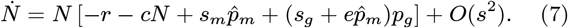

We are interested in the regime 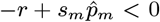, i.e. the microbe is not able to prevent extinction without spread of the rescue allele. The leading-order conditions for rescue follow immediately. The host allele alone eventually restores positive growth when *s*_*g*_ *> r*. With an endemic microbe the condition becomes

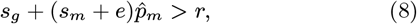

so the microbe is necessary for rescue when 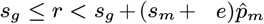.

For analytical interpretation, we retain only the leading-order forms in Eqs. 6–7 in the next section. Further below, we will discuss effects caused by higher order terms. For all maintext numerical trajectories, we substitute the first-order QME expressions,Eqs. 4–5, into the exact equations for *p*_*g*_ and *N* (Eqs. S1b and S2), without further expansion in selection. Thus, the reduced dynamics retain higher-order products generated by this substitution, but they do not constitute a systematic higher-order QME approximation.

### Additive and non-additive microbial effects shape rescue in distinct ways

Equations 6–7 separate the effects of the two microbial coefficients, *s*_*m*_ and *e*, cleanly. The additive coefficient *s*_*m*_ raises host growth by 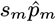 from the moment the environment deteriorates (Eq. 7), but it does not appear in the first-order equation for 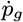, Eq. 6. It can thus lift the floor of the rescue-U (Fig. 2A) without touching the sweep rate (Fig. 2B); pure buffering (*s*_*m*_ *>* 0, *e* = 0) leaves the host allele trajectory exactly unchanged (with *e* = 0, the association 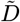 vanishes). The non-additive coefficient *e* does the opposite. It does not buffer initial decline (Fig. 2C), but enters 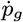 directly and, provided *e >* 0, accelerates the rescue-allele sweep (Fig. 2D), so its demographic effect is delayed but advances the sweep and shortens the excursion. Fig. 2E and F illustrate a scenario where both microbial effects are required for rescue. When *e <* 0 but *s*_*m*_ + *e >* 0, the microbe can remain beneficial to host growth (demographic buffering) while its effect on genetic adaptation is negative (evolutionary deceleration).

**Figure 2:**
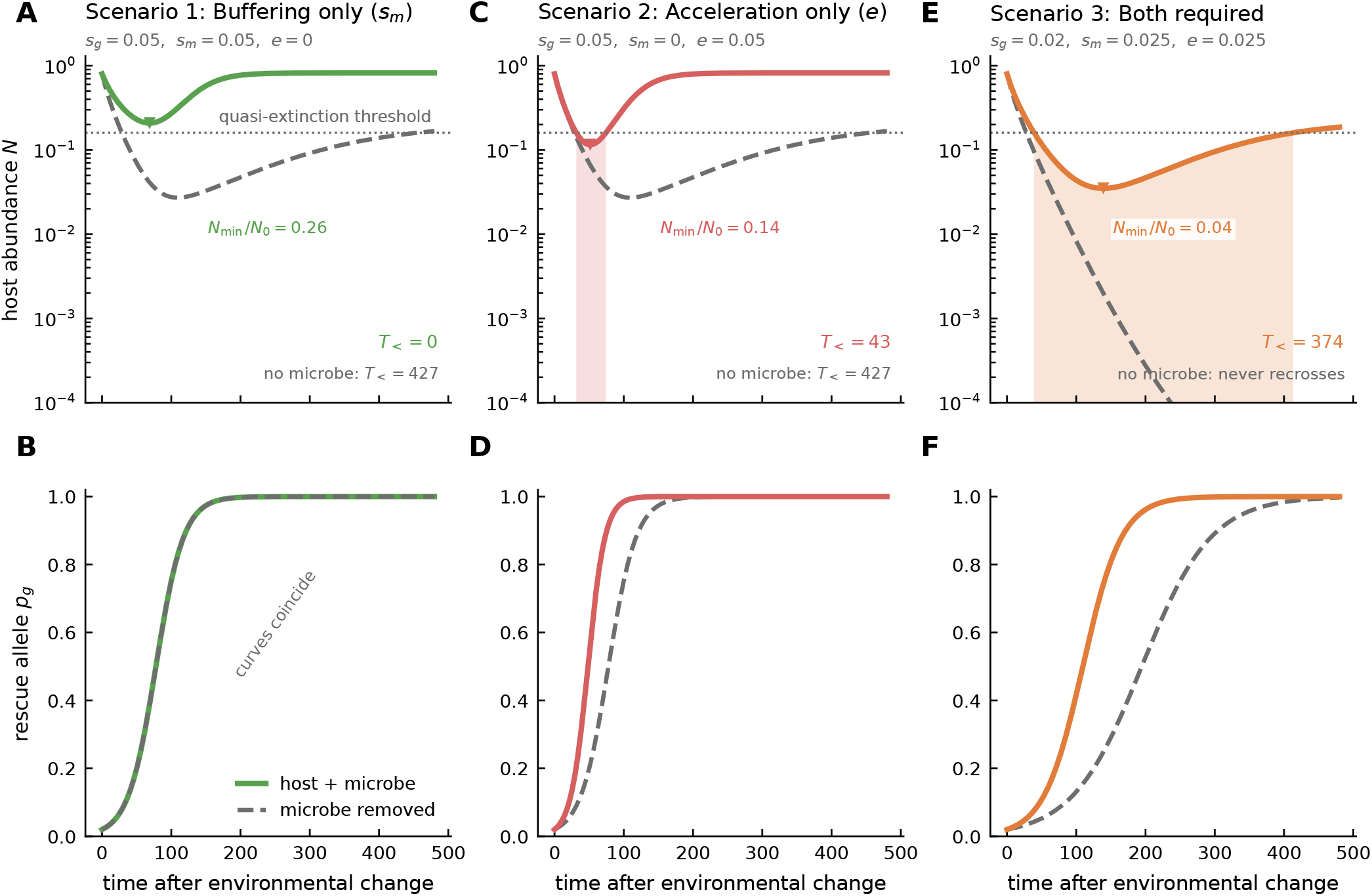
Three shapes of the rescue-U. Scenarios 1 and 2 hold *s*_*g*_ = 0.05 and the combined microbial coefficient *s*_*m*_ + *e* = 0.05 fixed, isolating whether microbial assistance acts through buffering or non-additivity; Scenario 3 lowers *s*_*g*_ so that both microbial routes are required. (*A,B*) Scenario 1: buffering only (*e* = 0) lifts the floor of the rescue-U while leaving the rescue-allele sweep exactly unchanged. (*C,D*) Scenario 2: acceleration only (*s*_*m*_ = 0) provides no allele-independent initial buffer but narrows the excursion by advancing the rescue-allele sweep. (*E,F*) Scenario 3: with *s*_*g*_ *< r* neither microbial route alone restores positive growth, but together they do. Dashed gray curves show results in the absence of the microbe. Shading marks time below an illustrative quasi-extinction threshold (dotted); *T*_*<*_ is the full duration from the first down-crossing to the next up-crossing, with trajectories that do not return labeled “never recrosses.” Full-model overlays are in SI Appendix, Fig. S1.

Figure 3 maps rescue-U depth and the time spent below an illustrative quasi-extinction threshold set to 20% of *N* (0) over *e* and *s*_*m*_, holding *s*_*g*_ *< r* so that the host allele cannot rescue on its own. The leading-order boundary 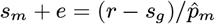 from Eq. 8 approximates the separation between positive and negative post-sweep low-density growth. Within the recovering region the two coefficients act on different features of the same excursion, so the same combined microbial coefficient produces visibly different rescue-U’s depending on how it is split. The white point marks the third scenario in Fig. 2.

**Figure 3:**
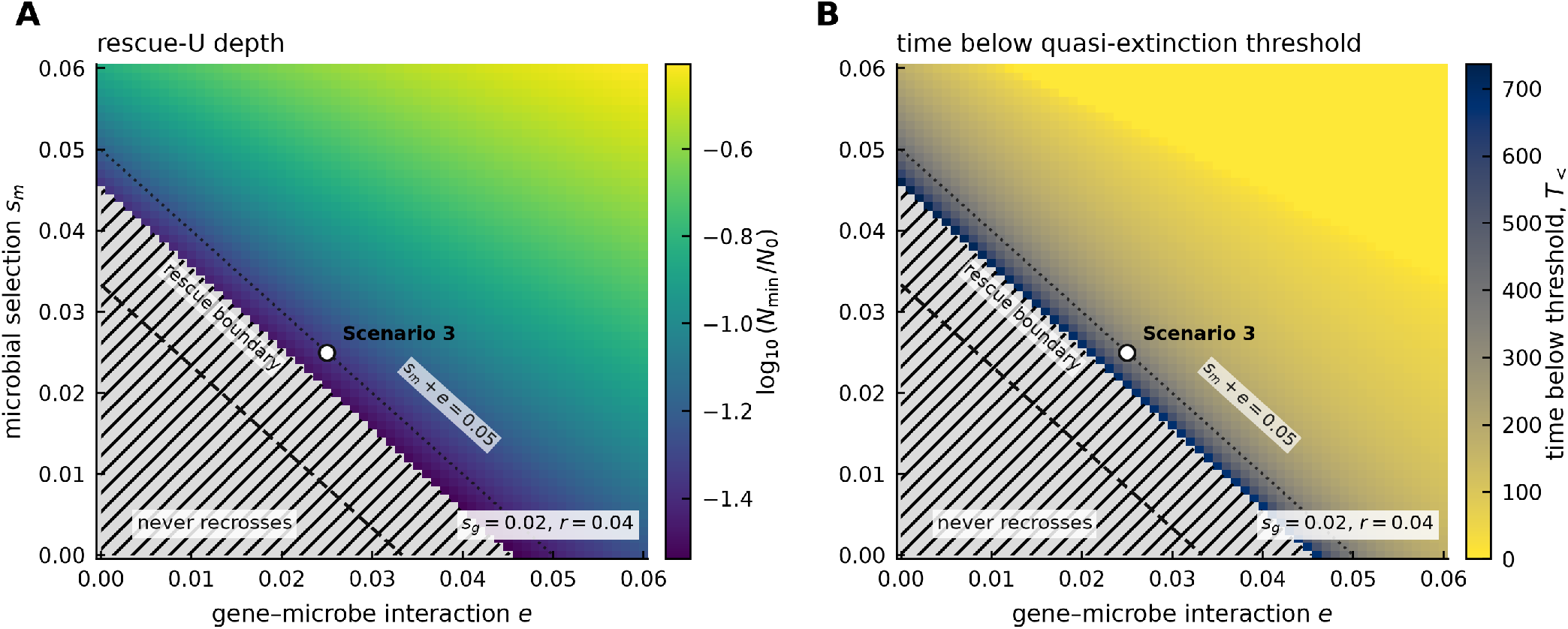
The two microbial coefficients shape different parts of the rescue-U. (*A,B*) Rescue-U depth and the full time below the quasi-extinction threshold over *e* and *s*_*m*_ at *s*_*g*_ = 0.02 *< r* = 0.04. Hatching marks parameter sets for which abundance never recrosses the threshold; only finite recrossing durations are colored in panel B. The dashed line is the leading-order rescue boundary 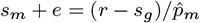 from Eq. (8), and the dotted line marks *s*_*m*_ + *e* = 0.05. The white point marks Scenario 3 in Fig. 2.

### Selection-amplified dilution can erase microbe-assisted rescue

Let *ϕ* = *f* (1 − *v*) denote fecundity-weighted failure to inherit the microbe, which generates the genotype-dependent microbial loss shown in Fig. 1A. Rescue is only possible if the low-density growth rate, once the rescue allele is fixed (*p*_*g*_ = 1), is larger than zero, i.e.:

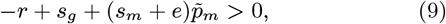

going now beyond the leading order in *s* (unlike in Eq. 8).

Equations 4–5 reveal that the response of microbial prevalence to the host allele sweep and the sign of host–microbe association are controlled by the same term:

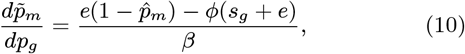

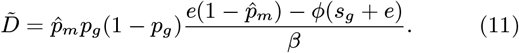

For *e >* 0, both change sign from positive to negative at

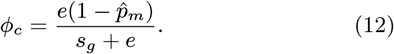

Hence, as the rescue allele sweeps to fixation (Fig. 4A), microbial prevalence increases or decreases, depending on Λ = *ϕ/ϕ*_*c*_: for Λ *<* 1, the beneficial host allele becomes positively associated with the microbe and microbial prevalence rises during the sweep; at Λ = 1, both responses vanish to first order; for Λ *>* 1, the host allele becomes negatively associated with the microbe and microbial prevalence falls as adaptation proceeds (Fig. 4B). Fig. 4C shows post-sweep low-density growth (scaled by *r*) as a function of Λ and 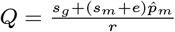 (the leading order rescue margin from Eq. (7)) and delineates regions in which rescue is or is not possible (separated by the solid black line) and in which microbial prevalence rises or falls (separated by the vertical dashed line at Λ = 1). A fall in microbial prevalence does not necessarily prevent the possibility of rescue (just as an increase does not guarantee it), but it can do so, as illustrated in Fig. 3D: for Λ = 0.5 and Λ = 1, the population size recovers eventually, while for Λ = 3, extinction is inevitable. Note that Λ also affects microbial prevalence at the beginning of the sweep but in the example, 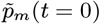 would have been large enough to allow for rescue even for Λ = 3.

**Figure 4:**
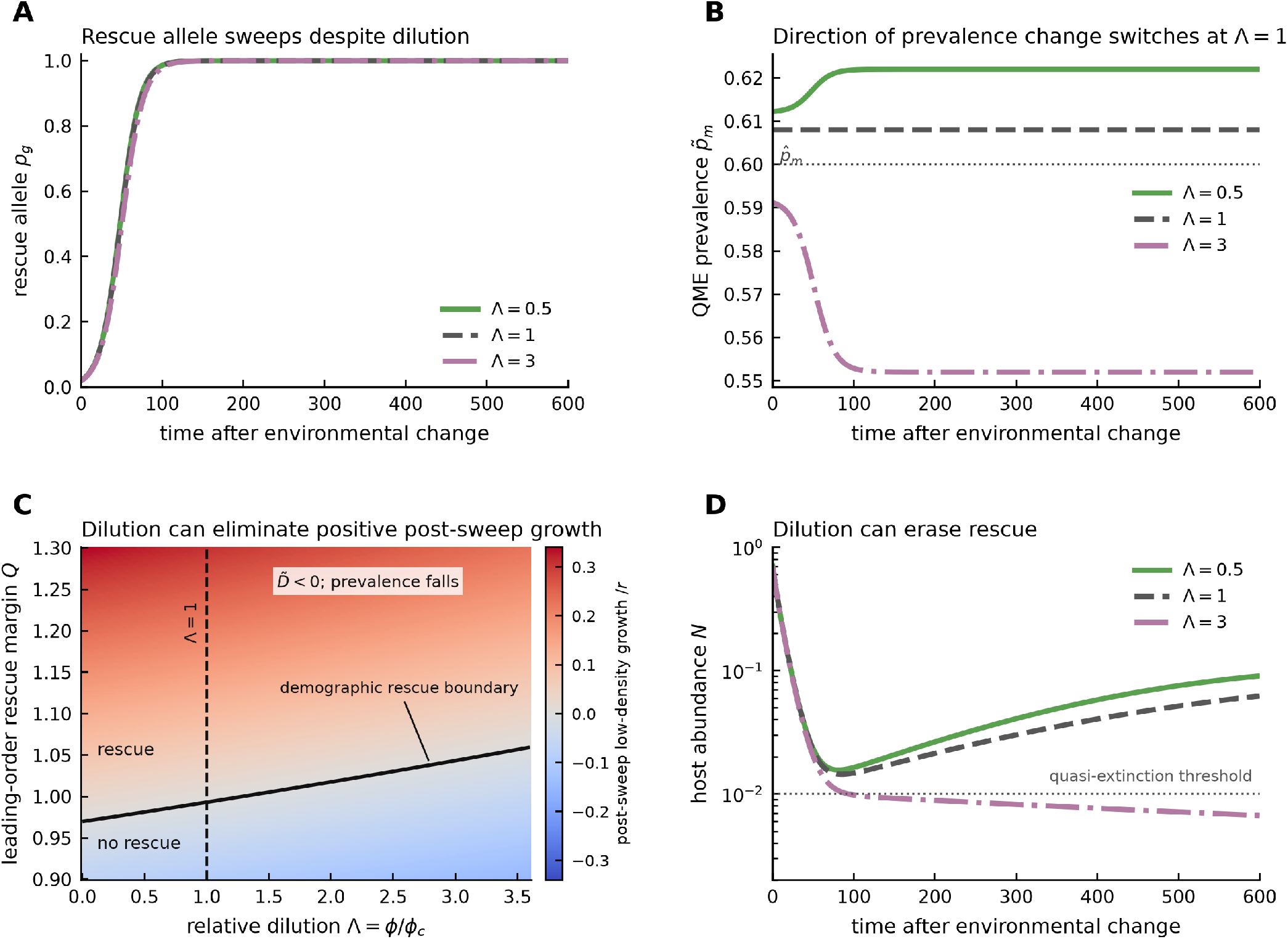
Selection-amplified dilution can erase rescue. (*A,B*) Representative QME trajectories show the rescue-allele sweep and the corresponding microbial prevalence: prevalence increases with the sweep for Λ *<* 1, is independent of *p*_*g*_ to first order at Λ = 1, and decreases with the sweep for Λ *>* 1, as predicted by Eqs. (10)–(12). (*C*) Post-sweep low-density growth across relative dilution and the leading-order rescue margin *Q*. The dashed line marks the prevalence/association threshold Λ = 1 from Eq. (12); the solid contour marks zero post-sweep low-density growth in the numerical QME reduction. (*D*) Corresponding abundance trajectories show recovery below and at the dilution threshold but failure of rescue under stronger dilution. Line types as well as colors distinguish Λ = 0.5, 1, and 3. Full-model overlays are in SI Appendix, Fig. S2. Representative trajectories use 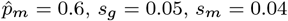, *e* = 0.05, and *r* = 0.10, giving *ϕ*_*c*_ = 0.20 and *Q* = 1.04.

Selection-amplified dilution belongs to the broader class of eco-evolutionary feedbacks in which individual-level adaptation does not optimize population persistence. Evolutionary-suicide theory shows that selection can drive populations toward lower abundance or across viability boundaries [21–23]. During environmental change, conflicts similar to selection-amplified dilution arise when selfing gains reproductive assurance or a transmission advantage while inbreeding depression lowers population growth [24, 25], and when cooperation is favored but its costs reduce effective population size and slow adaptation [26]. Here the selected host allele need not carry a direct demographic cost: its reproductive success instead increases the flux of descendants that fail to inherit a beneficial microbe. In the rescue setting analyzed here, this is most precisely an evolutionarily induced failure of rescue; if the same feedback drove an initially viable population across zero growth, it would constitute a microbial route to evolutionary suicide.

## Discussion

Microbes can alter evolutionary rescue by changing either side of the race between demographic decline and host adaptation. The additive microbial effect *s*_*m*_ slows demographic loss immediately, whereas the non-additive gene–microbe coefficient *e* changes how quickly the rescue allele spreads. A beneficial microbe can therefore buy time, accelerate adaptation, or do both, and these mechanisms leave different empirical signatures on time-series of host abundance and allele frequency dynamics. Additionally, a large effect of removing the microbiome (such as through axenic rearing) on host trait variation can coexist with little variation explained by naturally occurring microbiome composition. Furthermore, a microbe can alter a host trait without contributing independently transferable variation. Conversely, a modest average microbial effect can strongly alter adaptation when it is concentrated in particular host genotypes.

This separation connects several previous publications that have mostly developed in parallel. In detail, evolutionary-rescue theory identifies as the decisive quantities the initial decline rate, the rate of adaptive change, and the demographic threshold crossed during the lag [1, 27, 28]. In multilocus population genetics, non-additive fitness interactions and disequilibrium determine how fitness components combine and how rapidly associations are broken [18–20, 29]. In evolutionary epidemiology and symbiont theory, transmission, clearance, and vertical inheritance determine prevalence [3, 4]. The QME places these objects in one equation set. Microbial transmission–loss dynamics play the fast association-setting role that recombination plays in quasi-linkage equilibrium, but imperfect inheritance also couples selection to microbial loss. That extra term is what makes selection-amplified dilution possible.

The framework also connects directly to conjugative plasmids and antimicrobial resistance. In bacterial populations, plasmid carriage functions as an accessory locus: conjugation maps to horizontal acquisition, segregational loss to failed inheritance, and plasmid-borne resistance or metabolic genes to microbial fitness effects [30–32]. Plasmid-rescue models ask whether resistance establishes before antibiotic exposure eliminates the population and show that more horizontal transfer is not always better: resident plasmids can deplete recipients or exclude an incoming rescue plasmid [33–35]. Our results add the reciprocal prediction that adaptation of the chromosome or host background can itself erode a beneficial mobile element through selection-amplified dilution. The same transmission– selection conflict therefore spans host-associated microbes and plasmid-mediated antimicrobial resistance.

The selection-amplified dilution result broadens the usual expectation that beneficial microbes simply add resilience. The mechanism requires three ingredients: a fitness-enhancing host state, selection acting at least partly through reproduction, and imperfect transmission of the microbe through those additional births. It should be strongest when microbial prevalence is maintained by repeated acquisition rather than faithful vertical inheritance, and when host adaptation sharply increases fecundity. It should be weak when selection acts mainly through survival, when carriers transmit the microbe nearly perfectly, or when horizontal reacquisition is fast enough to refill the noncarrier class. These are measurable distinctions.

Three empirical systems map cleanly onto the model. In *Caenorhabditis elegans*, protective bacteria interact with host genotype and pathogen context, making factorial host genotype × microbe × pathogen experiments a direct route to *s*_*g*_, *s*_*m*_, and the non-additive interaction coefficient *e* [9, 36–39]. In *Daphnia*, clonal hosts, microbiome transplants, and population-level stress assays can separate average buffering from genotype-specific rescue and reveal interactions with other inherited symbionts [40]. In *Drosophila*, microbiome presence buffered an evolved developmental cost associated with insecticide resistance, whereas microbiota from resistant versus control populations did not transfer resistance-related phenotypes to naive hosts [41]; crossing resistant and control host backgrounds with axenic, conventional, and reciprocal-transplant treatments would test whether such buffering maps to transferable *s*_*m*_, a host-background-specific non-additive effect *e*, or neither. SI Appendix, Table S2 contains further details on experimental approaches that may be used to distinguish buffering, non-additive acceleration, and selection-amplified dilution. In every system, trait restoration matters for rescue only if it changes survival, fecundity, and population growth. Future research with these systems and others will be of particular value to further an empirical understanding of microbe-assisted rescue. Our model can generally guide the design of future empirical work. Across systems, the minimum informative dataset is a joint time series of host abundance, host genotype, microbial carriage or composition, and the life-history components generating growth. Endpoint sequencing alone cannot identify the mechanism: axenic comparisons estimate removal effects, reciprocal transplants estimate transferable microbial effects and host genotype interactions, and repeated sampling or pedigrees estimate persistence. Only their combination can distinguish buffering, acceleration, and dilution.

We anticipate that the most consequential extensions to our current model are biological rather than technical. Environmental reservoirs would create ecological memory: host shedding could maintain beneficial microbes between encounters, while spatially or temporally variable reacquisition could restore or erase host–microbe associations [42, 43]. Multiple strains would add competition, complementarity, cheaters, pathogens, transmission bottlenecks, and horizontal gene transfer, allowing rescue to depend on community composition rather than carriage of one element. Spatial structure could separate host and microbial expansion fronts, while coevolution of host filtering, microbial persistence, and transmission would turn *s*_*m*_, *e, β, v*, and *γ* into evolving quantities. Stochasticity remains essential at demographic bottlenecks, but these ecological extensions determine whether the single-microbe mechanisms scale to natural microbiomes.

Overall, our model integrates evolutionary rescue theory with multilocus population genetics and evolutionary epidemiology to enable a detailed characterization of how host-associated microbes can shape the demographic and evolutionary pathways that determine rescue. Our model framework highlights critical factors of the rescue process that warrant future empirical analysis. This framework additionally provides a backbone for further mathematical extensions that consider relevant ecological parameters likely shaping rescue in nature.

## Materials and Methods

### QME-reduced scenario analysis

All main-text trajectories were calculated from the QME-reduced system. For a given *p*_*g*_, Eqs. 4 and 5 supply 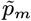 and 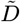. These QME values were substituted into the exact host allele equation

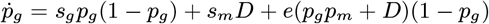

and into Eq. 2. This retains first-order changes in microbial prevalence and host–microbe association while avoiding direct integration of the fast microbial variables. For the host-only comparison, we remove the microbe by setting *p*_*m*_ = 0; consequently *D* = 0. Common competition was fixed within each comparison so differences in abundance arise from microbial and genetic contributions rather than altered density regulation.

For Fig. 1B–C, rates are expressed per host generation and we use *β* = 10, *ℓ*_0_ = 4, *s*_*g*_ = 0.03, *s*_*m*_ = 0.05, *e* = 0.10, and *ϕ* = 0, giving 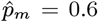. Panel B fixes *p*_*g*_ = 0.5 and starts from *p*_*m*_(0) = 0.55 and *D*(0) = 0, whereas panel C evaluates Eqs. 4–5 as *p*_*g*_ varies from 0 to 1. For Figs. 2–3, we set *β* = 1 to define the time unit; baseline prevalence is 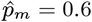, *r* = 0.04, *c* = 0.05, *ϕ* = 0, *p*_*g*_ (0) = 0.02, and *N* (0) = 0.8. The quasi-extinction threshold is set at 20% of *N* (0). We define *T*_*<*_ as the full duration from the first downward crossing to the next upward crossing; trajectories whose long-run density-regulated equilibrium remains below the threshold are labeled “never recrosses.” The rescue landscape varies *e* and *s*_*m*_ from 0 to 0.06 at *s*_*g*_ = 0.02. For Fig. 4, *s*_*g*_ = 0.05, *s*_*m*_ = 0.04, *e* = 0.05, 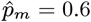, and *β* = 1; Λ varies from 0 to 3.6 and *Q* from 0.9 to 1.3. Representative dilution trajectories use *r* = 0.10, *c* = 0.05, *p*_*g*_ (0) = 0.02, and *N* (0) = 0.7.

### Numerical validation

The complete deterministic model was integrated for validation. Across the three rescue mechanisms, the maximum QME–full-model discrepancy in *p*_*g*_ is below 5.4 × 10^*−*4^ and the maximum discrepancy in log_10_ *N* below 3.6 × 10^*−*3^. In the strongest dilution trajectory (Λ = 3), where the approximation is more strongly stressed, the corresponding discrepancies are 3.6×10^*−*3^ and 0.082; both systems nevertheless predict failure of recovery. SI Appendix, Figs. S1–S2 show direct overlays. A broader numerical robustness analysis based on a 1,024-point Sobol design is given in SI Appendix, Fig. S3. The derivation and validation workflow is fully reproducible from the supplied Python scripts.

### Use of generative AI

Anthropic’s Claude Opus (versions 4.8–5) was used to assist in developing code for figure generation. OpenAI’s ChatGPT (GPT-5.5–5.6) was used for language editing and improving manuscript clarity. B.W. reviewed and validated the figure-generation code, verified the resulting figures, and reviewed all AI-assisted textual revisions. The authors take full responsibility for the final manuscript.

### Data, Materials, and Software Availability

No empirical data were generated. All equations are provided in the manuscript and Supporting Information. The Python scripts and parameter sets required to reproduce all main-text and Supporting Information figures are publicly available at https://github.com/bobweek/mash_figs.

## Author contributions

Conceptualization: B.W., H.U., and H.S.; methodology and formal analysis: B.W. and H.U.; software, validation, investigation, and visualization: B.W.; writing – original draft: B.W.; writing – review and editing: B.W., H.U., and H.S.; supervision: H.U. and H.S.; project administration: B.W. and H.S.; and senior authorship: H.U. and H.S. are co-senior authors.

## Acknowledgments

B.W. is funded through KiTE (Kiel Training for Excellence), which is part of the European Union’s Horizon Europe research and innovation programme under the Marie Sklodowska-Curie grant agreement number 101081480. H.S. is grateful for financial support from the German Science Foundation within the collaborative research centre CRC 1182 on the Origin and Function of Metaorganisms; project A4.3. H.S. further thanks the Max-Planck Society for a Max-Planck fellowship. We thank several participants of the workshop “Evolutionary rescue: bridging the gap between theory and experiments” for feedback on an earlier version of this project.

