## Supplementary Material for "Microbially assisted rescue of host populations"

### Supporting Information for
### Microbially assisted rescue of host populations

Bob Week, Hinrich Schulenburg, and Hildegard Uecker

#### S1. Full density-regulated host–microbe model

Let the four host types have frequencies $p_{\varnothing\varnothing}$, $p_{\varnothing m}$, $p_{g\varnothing}$, and $p_{gm}$, where $\varnothing$ denotes absence, $g$ presence of the focal host allele, and $m$ microbial carriage. These define the marginal frequencies

$$p_g = p_{g\varnothing} + p_{gm}, \qquad p_m = p_{\varnothing m} + p_{gm},$$

and the host–microbe association

$$D = p_{gm} - p_g p_m.$$

The selection coefficients are $s_m$ for microbial carriage and $s_g$ for the host allele; $e$ is the non-additive gene–microbe interaction coefficient. Horizontal acquisition is frequency dependent: each noncarrier acquires the microbe at rate $\beta p_m$, where $\beta$ is the transmission coefficient; baseline loss is $\ell_0 = b(1-v) + \gamma$, and $f(1-v)$ is the fraction of selection-dependent births that fail to inherit the microbe. Rewriting the four-class dynamics in terms of $(p_m, p_g, D)$ gives

$$\begin{aligned}
\dot{p}_m &= s_m p_m(1-p_m) + s_g D + e(p_m p_g + D)(1-p_m) \\
&\quad + \beta p_m(1-p_m) - \ell_0 p_m \\
&\quad - f(1-v)\left[s_m p_m + (s_g+e)(p_m p_g + D)\right], && \text{(S1a)}\\
\dot{p}_g &= s_g p_g(1-p_g) + s_m D + e(p_m p_g + D)(1-p_g), && \text{(S1b)}\\
\dot{D} &= e\left[p_m p_g(1-p_m)(1-p_g) + D(1-p_m-p_g) - D^2\right] \\
&\quad + D\left[s_m(1-2p_m) + s_g(1-2p_g) - (\beta p_m + \ell_0)\right] \\
&\quad - f(1-v)\left[(s_g+e)(1-p_g)(p_m p_g + D) + s_m D\right]. && \text{(S1c)}
\end{aligned}$$

Common competition enters only through abundance:

$$\dot{N} = N\left[-r - cN + s_g p_g + s_m p_m + e(p_g p_m + D)\right]. \qquad \text{(S2)}$$

Here $r > 0$, so $-r < 0$ is the baseline post-change Malthusian growth rate. Because $-cN$ is added equally to the Malthusian rate of every host class, it cancels exactly when class abundances are converted to frequencies. The original frequency model and its QME are therefore unchanged, while any positive low-density growth rate $\rho$ produces a finite ecological equilibrium $N^* = \rho/c$.

#### S2. Continuous-time Markov-chain and fluid-limit justification

Consider a population scaled by a system-size parameter $\Omega$, with counts

$$\mathbf{X}^\Omega = (X_{\varnothing\varnothing}, X_{\varnothing m}, X_{g\varnothing}, X_{gm})$$

and density vector $\mathbf{Z}^\Omega = \mathbf{X}^\Omega/\Omega$. Each individual gives birth at its state-specific fecundity, dies at its state-specific mortality plus common competitive mortality $cN$, and changes microbial state through acquisition or clearance. Carrier parents transmit the microbe with probability $v$; noninherited births enter the corresponding noncarrier class. Mendelian transmission of the haploid host allele is exact. Horizontal acquisition is frequency dependent, so each noncarrier changes state at rate $\beta p_m$. The finite process is therefore a density-dependent pure-jump Markov chain: every event changes $\mathbf{Z}^\Omega$ by $O(\Omega^{-1})$, while its total rate is $O(\Omega)$.

Writing the transition vectors as $\nu_k/\Omega$ and their density-dependent intensities as $\Omega q_k(\mathbf{z})$ gives

$$\mathbf{Z}^\Omega(t) = \mathbf{Z}^\Omega(0) + \sum_k \frac{\nu_k}{\Omega} Y_k\left(\Omega \int_0^t q_k(\mathbf{Z}^\Omega(u))\, du\right),$$

where the $Y_k$ are independent unit-rate Poisson processes. The law of large numbers for density-dependent Markov chains implies convergence on finite time intervals to

$$\dot{\mathbf{z}} = \sum_k \nu_k q_k(\mathbf{z})$$

as $\Omega \to \infty$ [1]. Transforming these four class-density equations to $(p_m, p_g, D, N)$ yields Eqs. S1–S2. The deterministic model is thus the fluid limit of an explicit birth–death–transmission process rather than a phenomenological frequency system.

This construction also identifies the natural stochastic extension. At finite $\Omega$, demographic and transmission noise are strongest near the rescue bottleneck. A diffusion approximation follows from the same event table by retaining the $O(\Omega^{-1/2})$ martingale term. For a final rescue analysis, the deterministic QME trajectory should provide the time-dependent drift, while branching-process or diffusion calculations determine the probability of crossing a quasi-extinction boundary.

### S3. QME derivation

Set the selective coefficients to a common weak scale,

$$s_g = \varepsilon a_g, \qquad s_m = \varepsilon a_m, \qquad e = \varepsilon a_e,$$

with $\varepsilon \ll 1$, while $\beta$ and $\ell_0$ remain $O(1)$. When $\beta > \ell_0$, the selection-free microbial subsystem has the stable endemic equilibrium

$$\hat{p}_m = 1 - \ell_0/\beta, \qquad D = 0.$$

We reserve hats for selection-free endemic equilibria and use tildes for quantities evaluated on the QME. We seek a slow manifold of the form

$$p_m = \hat{p}_m + \varepsilon u(p_g) + O(\varepsilon^2), \qquad D = \varepsilon d(p_g) + O(\varepsilon^2).$$

The host allele changes only at $O(\varepsilon)$, so $p_g$ is fixed when solving the $O(\varepsilon)$ fast-equilibrium conditions. Substituting the expansions into Eqs. S1a and S1c, setting their $O(\varepsilon)$ terms to zero, and then restoring the original coefficients gives

$$\tilde{p}_m = \hat{p}_m + \frac{(s_m + ep_g)(1-\hat{p}_m)}{\beta} - \frac{f(1-v)[s_m + (s_g+e)p_g]}{\beta} + O(s^2), \tag{S3}$$

$$\tilde{D} = \frac{\hat{p}_m p_g(1-p_g)}{\beta}\left[e(1-\hat{p}_m) - f(1-v)(s_g+e)\right] + O(s^2). \tag{S4}$$

Thus $\tilde{D}$ is the first-order host–microbe association on the QME; no separate approximation for the joint frequency is needed.

The relaxation eigenvalues of the selection-free fast subsystem are negative when the endemic state is stable. The QME is therefore justified when microbial transmission–loss dynamics relax substantially faster than host selection changes $p_g$. The approximation can remain qualitatively useful outside strict weak selection because its sign boundaries are often controlled by the same competing processes even when the first-order magnitudes are imperfect.

Substitution into the exact host allele equation yields

$$\dot{p}_g = (s_g + e\hat{p}_m)p_g(1-p_g) + O(s^2),$$

and substitution into Eq. S2 gives

$$\dot{N} = N\left[-r - cN + s_m\hat{p}_m + (s_g + e\hat{p}_m)p_g\right] + O(s^2).$$

These expressions establish the first-order separation between microbial buffering and non-additive alteration of host adaptation.

##### Selection-amplified dilution boundaries

Let $\phi = f(1-v)$. Differentiating Eq. S3 with respect to the slow host allele frequency gives

$$\frac{d\tilde{p}_m}{dp_g} = \frac{e(1-\hat{p}_m) - \phi(s_g + e)}{\beta} + O(s^2). \tag{S5}$$

The same factor multiplies the first-order association,

$$\tilde{D} = \frac{\hat{p}_m p_g(1-p_g)}{\beta}\left[e(1-\hat{p}_m) - \phi(s_g+e)\right] + O(s^2). \tag{S6}$$

For $e > 0$, the sign of microbial change along the host sweep is determined by the same threshold as the sign of the association:

$$\phi_c = \frac{e(1-\hat{p}_m)}{s_g + e}. \tag{S7}$$

The dimensionless ratio $\Lambda = \phi/\phi_c$ is used in main-text Fig. 4. When $\Lambda < 1$, $\tilde{p}_m$ increases with $p_g$ and $\tilde{D} > 0$ for $0 < p_g < 1$; at $\Lambda = 1$ both responses vanish to first order; when $\Lambda > 1$, $\tilde{p}_m$ decreases as the beneficial host allele rises and $\tilde{D} < 0$. The microbial selection coefficient $s_m$ does not enter this sign boundary because it does not distinguish host allele backgrounds at first order.

#### S4. Rescue scenarios, parameter sweeps, and validation

At leading order the host allele sweeps at rate $s_g + e\hat{p}_m$ and low-density growth is $-r + s_m\hat{p}_m + (s_g + e\hat{p}_m)p_g$. Host-only rescue therefore requires $s_g > r$, whereas an endemic microbe restores positive post-sweep low-density growth when $s_g + (s_m + e)\hat{p}_m > r$. The main-text trajectories were selected to isolate the two microbial coefficients rather than to maximize visual separation. All three hold the combined microbial coefficient $s_m + e$ fixed; the first two also hold $s_g$ fixed, whereas the third lowers $s_g$ to create a microbe-required regime. Table S1 lists the parameters.

Table S1: Parameter values for main-text trajectory scenarios. All rescue scenarios use $\beta = 1$, $\ell_0 = 0.4$ ($\hat{p}_m = 0.6$), $r = 0.04$, $c = 0.05$, and $\phi = f(1-v) = 0$, and all hold the combined microbial coefficient $s_m + e = 0.05$. The final column reports the leading-order post-sweep low-density growth rate $s_g + (s_m + e)\hat{p}_m - r$, which is positive exactly when the population recovers.

| Scenario | $s_g$ | $s_m$ | $e$ | $s_m + e$ | post-sweep low-density growth |
|---|---|---|---|---|---|
| Buffering only | 0.050 | 0.050 | 0 | 0.05 | +0.040 |
| Acceleration only | 0.050 | 0 | 0.050 | 0.05 | +0.040 |
| Both required | 0.020 | 0.025 | 0.025 | 0.05 | +0.010 |

The pure-buffering scenario is exactly useful as a control: because $e = 0$ and $\phi = 0$, the microbe does not alter $p_g$ at the order retained, so the host-only and host-plus-microbe allele trajectories overlap while abundance differs. In the pure-acceleration scenario $s_m = 0$, so there is no initial additive microbial buffer; the difference arises from the non-additive interaction. In the microbe-required scenario $s_g = 0.02 < r$, and $s_g + s_m\hat{p}_m = s_g + e\hat{p}_m = 0.035 < r = 0.04$ while $s_g + (s_m + e)\hat{p}_m = 0.05 > r$, so neither coefficient alone restores positive post-sweep low-density growth at leading order but their combination does. For the illustrative threshold in Fig. 2, the microbial trajectories have $T_< = 0$, 43, and 374 time units for buffering, acceleration, and the both-required scenario, respectively. The corresponding microbe-free trajectory has $T_< \approx 427$ time units when $s_g = 0.05$ and never recrosses the threshold when $s_g = 0.02$.

For the rescue maps in main-text Fig. 3, we fixed $\hat{p}_m = 0.6$, $\beta = 1$, $r = 0.04$, $c = 0.05$, $p_g(0) = 0.02$, $N(0) = 0.8$, and $\phi = 0$, and varied $e, s_m \in [0, 0.06]$ on a regular grid at $s_g = 0.02$. The plotted outputs are the minimum relative abundance $N_{\min}/N_0$ and $T_<$, defined as the full duration from the first downward crossing of a quasi-extinction threshold set at 20% of $N(0)$ to the next upward crossing. The numerical integration is continued until every trajectory with a long-run equilibrium above the threshold has recrossed it; parameter sets whose density-regulated equilibrium remains below the threshold are instead marked "never recrosses." The threshold is illustrative: it is a reference level for comparing excursions, not a claim about any particular extinction risk. The leading rescue boundary is the straight line $s_m + e = (r - s_g)/\hat{p}_m$. The numerical QME reduction shifts this boundary only slightly for the weak coefficients used here.

The full system integrates Eqs. S1–S2 from the same initial point on the QME manifold. Maximum absolute QME–full discrepancies over the plotted interval $(p_g, p_m, D, \log_{10} N)$ were $(< 10^{-12}, 9.5 \times 10^{-4}, 0, 1.2 \times 10^{-3})$ for pure buffering, $(2.5 \times 10^{-4}, 9.5 \times 10^{-4}, 7.2 \times 10^{-5}, 1.0 \times 10^{-3})$ for pure acceleration, and $(5.3 \times 10^{-4}, 9.5 \times 10^{-4}, 5.3 \times 10^{-5}, 3.6 \times 10^{-3})$ when both routes are required. The QME therefore reproduces both the evolutionary timing and bottleneck depth in all three representative regimes.

### S5. Selection-amplified dilution map and validation

Main-text Fig. 4 uses $\beta = 1$, $\hat{p}_m = 0.6$, $s_g = 0.05$, $s_m = 0.04$, and $e = 0.05$, which gives $\phi_c = 0.20$. The representative trajectories set $r = 0.10$, so the no-dilution rescue margin is

$$Q = \frac{s_g + (s_m + e)\hat{p}_m}{r} = 1.04.$$

They compare $\Lambda = 0.5, 1$, and 3, corresponding to $\phi = 0.10, 0.20$, and 0.60. The first two trajectories recover deterministically; the last has negative post-sweep low-density growth and continues to decline. The dotted abundance line in the main figure is an illustrative quasi-extinction threshold and is not used to define deterministic rescue.

For the phase map, $\Lambda$ varies from 0 to 3.6 and $Q$ from 0.9 to 1.3. We vary $Q$ by changing $r$ while holding the selective and transmission parameters fixed. Colors show post-sweep low-density growth in the numerical QME reduction divided by $r$. At $p_g = 1$, $\tilde{D} = 0$, so the demographic rescue boundary satisfies $-r + s_g + (s_m + e)\tilde{p}_m(1) = 0$. The vertical line $\Lambda = 1$ is the analytic prevalence/association threshold from Eq. S7. Because the numerical reduction substitutes the first-order QME into the exact growth equation, its zero-growth contour differs slightly from the leading-order reference $Q = 1$ even at zero dilution.

Thus $\Lambda = 0$, not $\Lambda = 1$, represents absence of dilution. The threshold $\Lambda = 1$ instead concerns the direction of prevalence change during the host sweep and the sign of association; it need not coincide with the demographic condition $Q = 1$. For the parameters of Fig. 4, the numerical-QME demographic boundary crosses $Q = 1$ at $\Lambda \approx 1.29$, where $\tilde{p}_m(1) = \hat{p}_m$.

For $\Lambda = 0.5, 1$, and 3, maximum absolute QME–full discrepancies over the plotted interval $(p_g, p_m, D, \log_{10} N)$ were respectively $(9.9 \times 10^{-4}, 1.8 \times 10^{-3}, 1.4 \times 10^{-4}, 0.020)$, $(4.6 \times 10^{-4}, 6.6 \times 10^{-4}, 5.5 \times 10^{-5}, 0.0086)$, and $(3.5 \times 10^{-3}, 4.0 \times 10^{-3}, 6.4 \times 10^{-4}, 0.081)$. Quantitative error grows in the strongest-dilution case, but the full model retains decreasing microbial prevalence, negative association, and failure of demographic recovery.

### S6. Broader numerical validation of the QME

The original robustness analysis compared four sign predictions: the selection-induced deviation $\Delta p_m = p_m - \hat{p}_m$; the sign of the association $D$; the microbial contribution to effective selection on the focal host allele,

$$A_m = \frac{\dot{p}_g}{p_g(1-p_g)} - s_g = e\left(p_m + \frac{D}{p_g}\right) + \frac{s_m D}{p_g(1-p_g)};$$

and the microbial contribution to growth,

$$G_m = s_m p_m + e(p_g p_m + D).$$

At first order the corresponding QME quantities are $\tilde{p}_m - \hat{p}_m$, $\tilde{D}$, $e\hat{p}_m$, and $\hat{p}_m(s_m + e p_g)$.

We evaluated $2^{10} = 1,024$ parameter sets generated with a scrambled Sobol low-discrepancy design [2, 3], implemented in SciPy. With $\beta = 1$ as the rate scale, the largest absolute selective coefficient ranged from approximately 0.005 to 0.5. The coefficients $s_m$, $s_g$, and $e$ varied independently in sign; $\hat{p}_m$ ranged from 0.05 to 0.95 and $f(1-v)$ from 0 to 0.8. Full nonlinear conditional equilibria supplied initial $p_m$ and $D$ at $p_g = 0.1$. Biologically admissible trajectories were obtained for 1,005 sets.

Away from zero boundaries, the QME sign was retained for the entire full-model trajectory in 99.0% of resolved cases for microbial prevalence, 98.7% for association, 99.3% for the microbial effect on host adaptation, and 99.2% for the microbial contribution to growth. The sign of total effective selection on the focal host allele was retained in 99.7% of resolved trajectories. Even when the largest selective coefficient was approximately 0.23–0.5 times the transmission rate, agreement remained between 93% and 96% for the four microbial-effect signs (Fig. S3). Sign reversals were concentrated near QME zero boundaries, where higher-order terms are expected to dominate.

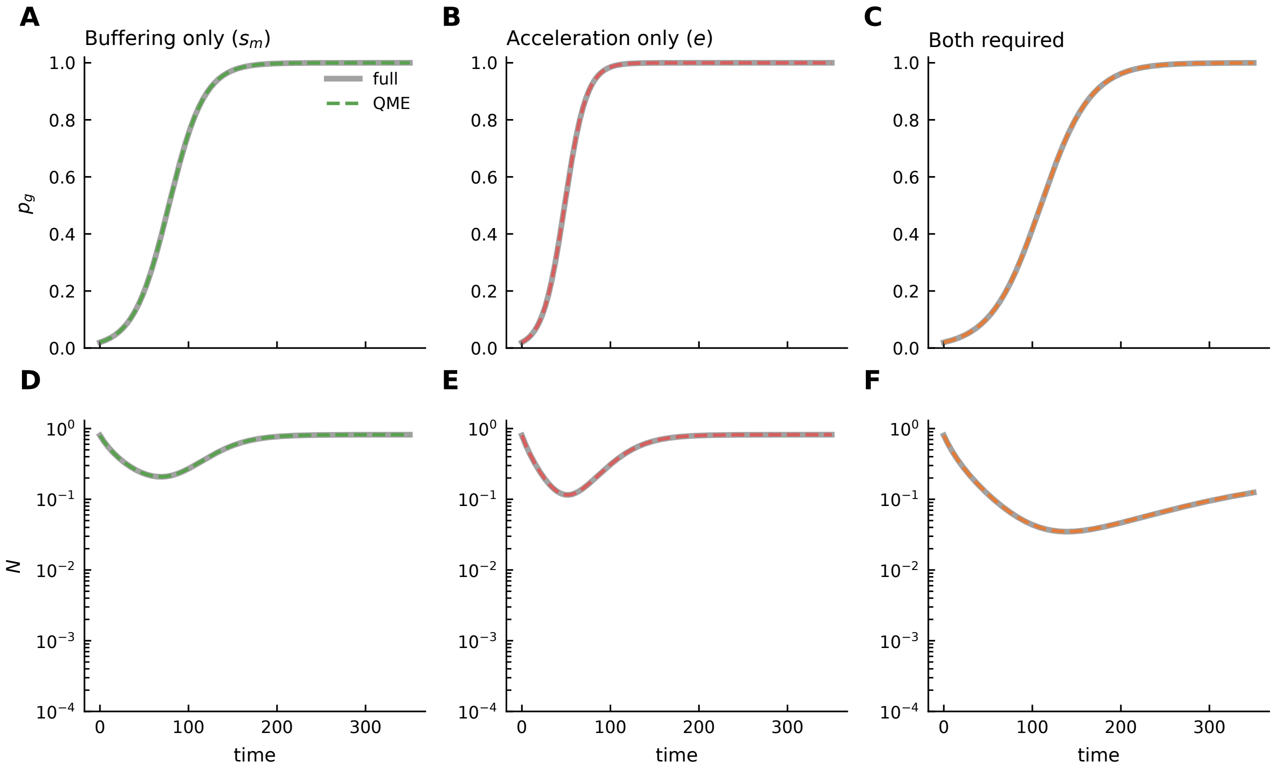


Figure S1: **Full-model validation of the three rescue mechanisms.** Solid gray curves are the full density-regulated model; dashed colored curves are the QME reduction. Columns correspond to pure buffering, pure non-additive acceleration, and the case in which both are required for rescue.

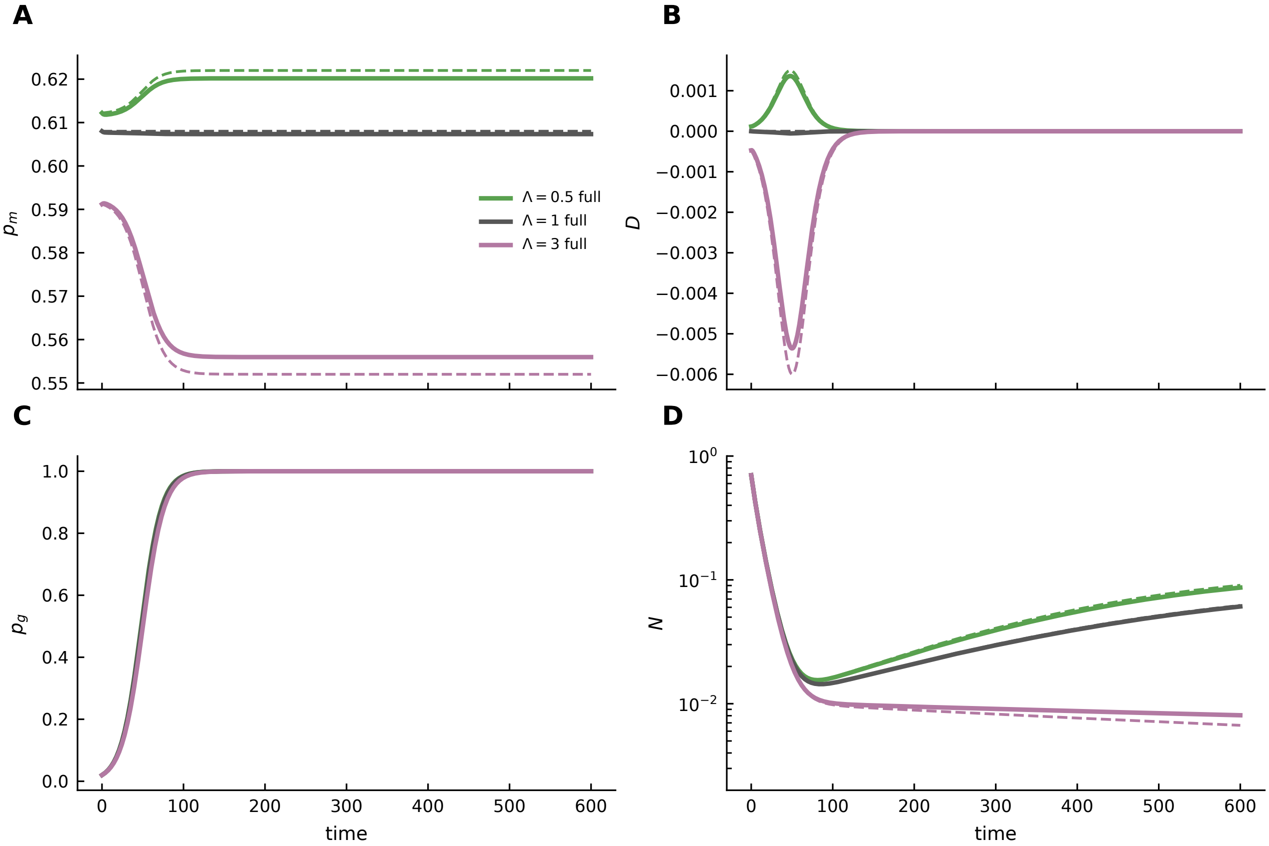


Figure S2: **Full-model validation of the dilution threshold and rescue failure.** Solid curves show the full model; dashed curves show the QME. Colors distinguish $\Lambda = 0.5, 1$, and 3. The QME captures microbial prevalence (A), association (B), the host sweep (C), and abundance (D), including continued decline at $\Lambda = 3$.

### S7. Empirical systems and measurements

The main-text discussion highlights three systems in which the model parameters can in principle be separated experimentally. Table S2 below proposes further details on what experiments would be possible with each system, and what quantities may be identified by such experiments.

Table S2: Empirical measurements that would distinguish microbial buffering, non-additive acceleration, and selection-amplified dilution.

| System | Existing empirical basis | Critical experiment | Quantities identified |
|---|---|---|---|
| *C. elegans*–protective bacteria–pathogen | Defined host genotypes, cultured protective strains, tractable infection and adaptation assays [4–8] | Cross host genotype × microbe presence / strain × stress; follow carriage through reproduction, clearance, and reacquisition | $s_g$, $s_m$, $e$ on survival/fecundity; $\beta$, $v$, $\gamma$; sign of $D$ |
| *Daphnia* under environmental stress | Clonal hosts, microbiome transplants, population-level stress assays [9] | Reciprocal microbiome inocula across clones with host-abundance time series and joint symbiont / pathogen profiling | Buffering versus genotype-specific effects; rescue threshold; interactions among symbionts |
| Evolved and control *Drosophila* | Host adaptation, axenic manipulation showing microbiome-dependent buffering of development, and resistant-versus-control microbiome transplants with no detectable host-phenotype effect [10] | Factorial host background × axenic / conventional / transplant treatment; measure development, survival, fecundity, and carriage jointly | Transferable $s_m$ versus genotype-specific $e$; whether trait restoration changes total growth |

### S8. Reproducibility

All code required to reproduce the numerical figures is publicly available at `https://github.com/bobweek/mash_figs`. The script `scripts/make_pnas_figures.py` reproduces main-text Figs. 1–4 and SI Figs. S1–S3. The module `scripts/qme_rescue_model.py` contains the full frequency model, QME functions, analytic dilution threshold, physical-state checks, numerical solvers, and the utilities used for the Sobol robustness analysis. Together these two files reproduce all numerical figures used in the manuscript. Running the figure-generation script from the repository root automatically creates a `figures/` directory containing the figures in PDF and PNG formats.

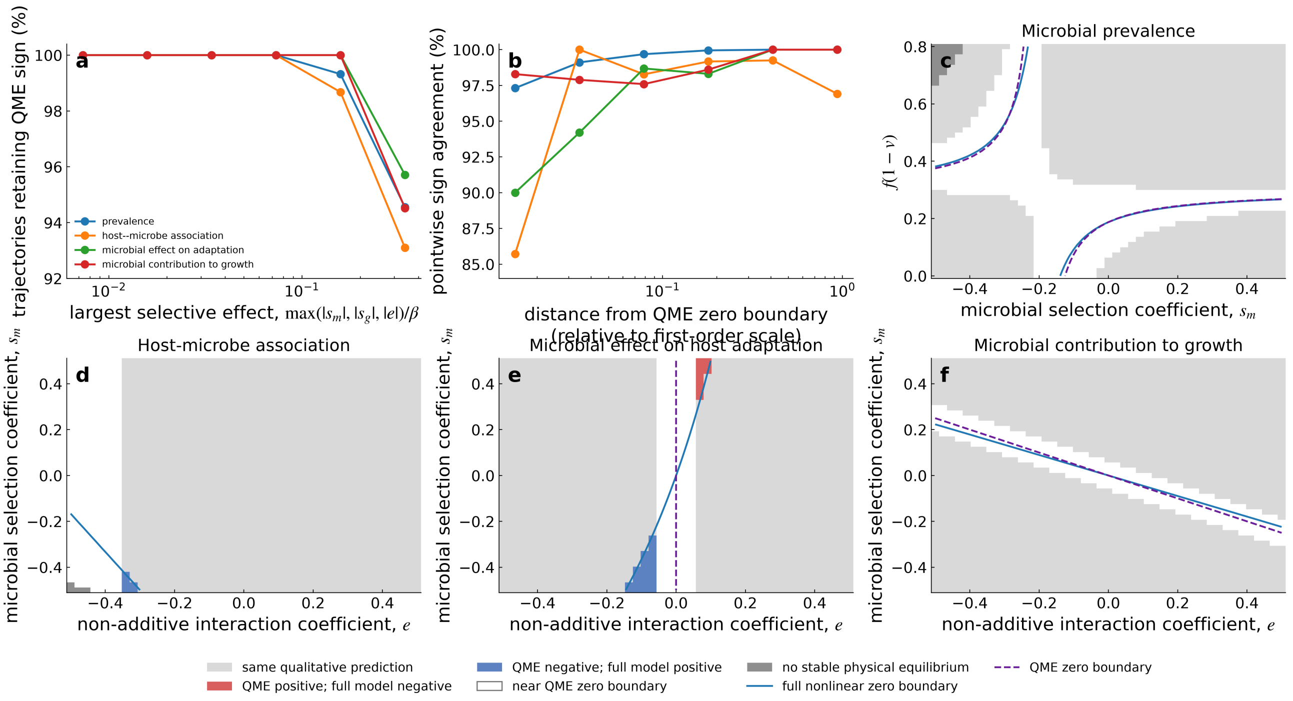


Figure S3: **QME sign predictions remain robust beyond strict weak selection.** Panels summarize trajectory-wide and pointwise sign agreement across the Sobol design and compare full nonlinear with QME zero boundaries in representative strong-selection slices.
